# Assessing the reliability of spike-in normalization for analyses of single-cell RNA sequencing data

**DOI:** 10.1101/119784

**Authors:** Aaron T. L. Lun, Fernando J. Calero-Nieto, Liora Haim-Vilmovsky, Berthold Göttgens, John C. Marioni

**Affiliations:** Cancer Research UK Cambridge Institute, University of Cambridge, Li Ka Shing Centre, Robinson Way, Cambridge CB2 0RE, United Kingdom; Wellcome Trust and MRC Cambridge Stem Cell Institute, University of Cambridge, Wellcome Trust/MRC Building, Hills Road, Cambridge CB2 0XY, United Kingdom; EMBL European Bioinformatics Institute, Wellcome Genome Campus, Hinxton, Cambridge CB10 1SD, United Kingdom; Wellcome Trust Sanger Institute, Wellcome Genome Campus, Hinxton, Cambridge CB10 1SA, United Kingdom

## Abstract

By profiling the transcriptomes of individual cells, single-cell RNA sequencing provides unparalleled resolution to study cellular heterogeneity. However, this comes at the cost of high technical noise, including cell-specific biases in capture efficiency and library generation. One strategy for removing these biases is to add a constant amount of spike-in RNA to each cell, and to scale the observed expression values so that the coverage of spike-in RNA is constant across cells. This approach has previously been criticized as its accuracy depends on the precise addition of spike-in RNA to each sample, and on similarities in behaviour (e.g., capture efficiency) between the spike-in and endogenous transcripts. Here, we perform mixture experiments using two different sets of spike-in RNA to quantify the variance in the amount of spike-in RNA added to each well in a plate-based protocol. We also obtain an upper bound on the variance due to differences in behaviour between the two spike-in sets. We demonstrate that both factors are small contributors to the total technical variance and have only minor effects on downstream analyses such as detection of highly variable genes and clustering. Our results suggest that spike-in normalization is reliable enough for routine use in single-cell RNA sequencing data analyses.

## Introduction

Single-cell RNA sequencing (scRNA-seq) is a powerful technique for studying transcriptional activity in individual cells. Briefly, RNA is isolated from single cells, reverse transcribed into cDNA and sequenced using massively parallel sequencing technologies [28]. This can be performed using microfluidics platforms like the Fluidigm C1 [21]; with protocols such as Smart-seq2 [20] that use microtiter plates; or with droplet-based technologies [12, 17] that can profile thousands of cells. Gene expression is 7 quantified by mapping read sequences to a reference genome and counting the number of reads mapped to each annotated gene. To avoid amplification biases, individual transcript molecules can also be tagged with unique molecular identifiers (UMIs) [10], such that sequencing to saturation and counting UMIs will yield the number of transcripts of each gene in a cell. Regardless of whether reads or UMIs are used, not all transcript molecules will be captured and sequenced due to cell-specific inefficiencies in reverse transcription [29]. The presence of these cell-specific biases compromises the direct use of the read/UMI count as a quantitative measure of gene expression. Normalization is required to remove these biases before the gene counts can be meaningfully compared between cells in downstream analyses.

A common normalization strategy for RNA-seq data uses a set of genes that have constant expression across cells. This set can consist of pre-defined “house-keeping” genes, or it can be empirically defined under the assumption that most genes are not differentially expressed (DE) between cells [1, 15, 24]. Any systematic differences in expression between cells for this non-DE set of genes must, therefore, be technical in origin, e.g., due to differences in library size or composition bias [24]. Counts are scaled to eliminate these differences, yielding normalized expression values for downstream analyses. This gene-based approach works well for bulk sequencing experiments where the population-wide gene expression profile is stable. However, it may not be suitable for single-cell experiments where strong biological heterogeneity complicates the identification of a reliable non-DE set. For example, house-keeping genes may be turned on or off by transcriptional bursting, while processes like the cell cycle may trigger large-scale changes in the expression profile that preclude a non-DE majority.

An alternative normalization approach is to use spike-in RNA for which the identity and quantity of all transcripts is known [2, 29]. The same amount of spike-in RNA is added to each cell’s lysate, and the spike-in transcripts are processed in parallel with their endogenous counterparts to generate a sequencing library. This yields a set of read (or UMI) counts for both endogenous and spike-in transcripts in each cell. Normalization is performed by scaling the counts for each cell such that the counts for the spike-in genes are, on average, the same between cells [11]. The central assumptions of this approach are that (i) the same amount of spike-in RNA is added to each cell, and (ii) the spike-in and endogenous transcripts are similarly affected by cell-to-cell fluctuations in capture efficiency. Under these assumptions, any differences in the coverage of the spike-in transcripts between cells must be artifactual in origin and should be removed by scaling. One particular advantage of this strategy is that it does not make any assumptions about the endogenous expression profile, unlike the non-DE approach described above. This means that spike-in normalization can be applied in situations where large-scale changes in expression (e.g., related to changes in total RNA content, or involving highly heterogeneous populations containing many cell types) are expected and of interest [16, 19].

There are two common criticisms of spike-in normalization that challenge the validity of its central assumptions. The first is that the same quantity of spike-in RNA may not be consistently added to each sample [24], and the second is that synthetic spike-in transcripts may not behave in the same manner as endogenous transcripts [6] (i.e., unequal capture efficiencies, caused by differences in the biophysical properties of the transcripts). Any differences in spike-in quantity or behaviour across cells will compromise the accuracy of spike-in normalization [22]. In some cases, it may also be difficult to gauge how much spike-in RNA should be added, especially if the quantity of endogenous RNA per cell is unknown, resulting in insufficient spike-in coverage for normalization. These criticisms may contribute to the limited use of this normalization strategy in the scRNA-seq literature [2]. However, if one were to dismiss the use of spike-in normalization, there would be no general alternative for removing cell-specific biases in scRNA-seq data sets where a non-DE majority of genes cannot be assumed. Thus, it is of particular interest whether or not the aforementioned criticisms of spike-in normalization are relevant to real scRNA-seq experiments. To our knowledge, this has yet to be rigorously studied.

In this paper, we conduct a series of experiments to estimate the reliability of spike-in normalization in single-cell transcriptome studies employing plate-based protocols. We use mixtures of two distinct spike-in RNA sets to quantify the variance of the added spike-in volume across cells, and show that it is quantitatively negligible in real experiments across a range of conditions. We also obtain an upper bound on the cell-to-cell variability in the differences in behaviour (i.e., the fold-changes in the capture efficiencies) between the two spike-in sets. Simulations indicate that both factors have only minor effects on the results of downstream analyses such as detection of DE and highly variable genes. These results suggest that spike-ins can be safely used for routine normalization of scRNA-seq data.

## Results

### Overview of the mixture experiments

We aimed to assess the variability in the added spike-in quantity across cells. To do so, we performed mixture experiments using two distinct spike-in sets (Figure 1) – the External RNA Controls Consortium (ERCC) set and the Spike-in RNA Variants (SIRV) set. An equal volume of each spike-in set was added separately to all wells of a 96-well microtiter plate. Each well contained a single lysed mouse cell – a mouse 416B myeloid progenitor cell or trophoblast stem cell (TSC) – thus mimicking real experimental conditions. The resulting pool of endogenous/spike-in RNA in each well was used to generate a cDNA library, using a modified version of the Smart-seq2 protocol (see Methods). This process was repeated for all wells and high-throughput sequencing was performed on all libraries.

**Figure 1.**
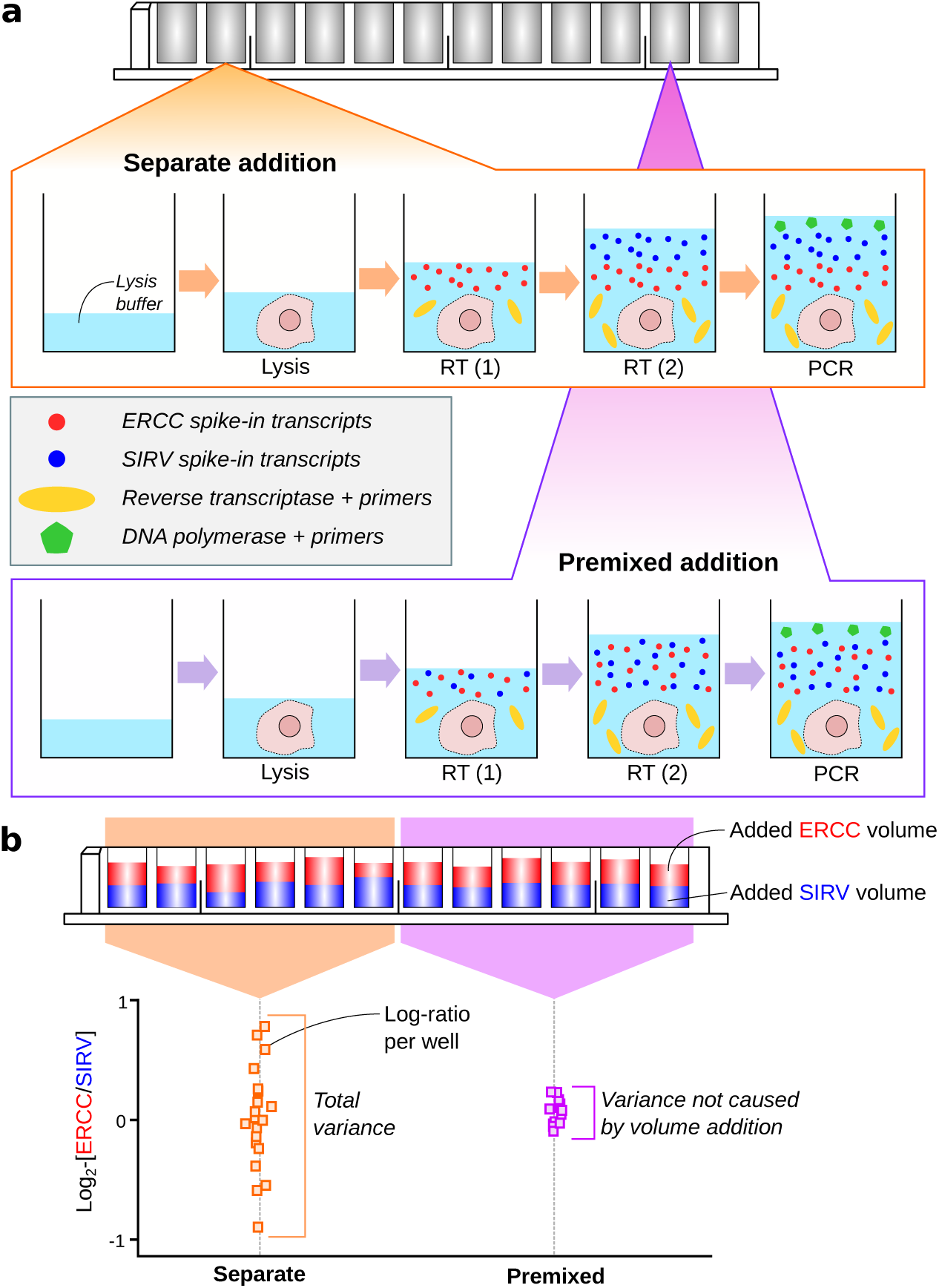
Schematic of the experimental design to assess the variability of spike-in addition in a plate-based scRNA-seq protocol. (a) A cell is sorted into each well of a plate and lysed. For one set of wells, an equal volume of each spike-in set is added separately, along with the reverse transcription (RT) reagents. For another set of wells, an equal volume of a pooled mixture of the two spike-ins is added into each well (done twice to keep the protocol consistent). Reverse transcription, PCR amplification, library generation and sequencing were then performed. (b) The log_2_-ratio between the total counts of the two spike-in sets was computed for each well. The variance of the log-ratio was estimated from all wells with separate addition of spike-ins, and from wells with addition of the premixed pool. The difference between these two estimates represents the variance attributable to volume addition.

For each library, reads were mapped to the genome and assigned to genes to quantify expression. The total count was computed across all transcripts of each spike-in set in each well. The log_2_-ratio of the totals between the two sets was computed for each well, and the variance of this log-ratio was computed *across* wells. Any variability in spike-in volume addition should manifest as an increase to the variability of the log-ratio, given that the spike-in sets were added independently to each well.

We also repeated the experiment by adding volumes of “premixed” spike-in solution where the two spike-in sets had been pooled at a 1:1 ratio. This ensures that there is no well-to-well variability in the relative quantities of RNA from the two spike-in sets. The variance of the log-ratio across these premixed-addition wells provides a baseline level of variability in the protocol (e.g., due to sequencing noise). The variance of volume addition was then estimated as the difference in the variance estimates from the premixed-addition wells and from the wells with separate addition of spike-ins.

We performed both the premixed and separate-addition experiments on the same plate to avoid plate effects [8, 30]. For the separate-addition experiment, we also reversed the order of addition of the two spike-in sets to determine if this affected the variance estimate. Finally, we generated data from replicate plates to ensure our results were reproducible. This was done in a range of conditions, i.e., using different cell types, by different operators and with sequencing at different locations.

We used a protocol based on microtiter plates rather than microfluidics as it is easier to customise the spike-in addition step in the former. Our experimental design requires two separate additions of spike-in RNA to each reaction (see Methods). This is not straightforward to achieve on, say, the Fluidigm C1 chip where the added volume for each reagent depends on the design on the reaction chamber. Our focus on data from plate-based protocols reflects their widespread use in single-cell studies [9, 26, 27, 32]. Obviously, the procedure we describe here can be adapted to any protocol where the spike-in addition can be easily modified, e.g., plate-based CEL-seq [7] or STRT-seq [9].

### Estimating the variance of volume addition

Denote the log_2_-transformed total read count for well *i* and spike-in set *s* as

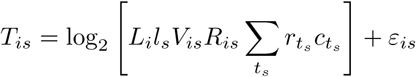
where the sum is taken over all unique transcripts *t_s_* in *s*. The other terms are defined as follows:

- *c_t_s__*, a constant specifying the concentration (in terms of transcripts per unit of volume) of *t_s_*.
- *r_t_s__*, a constant specifying the optimal transcript molecule-to-cDNA fragment capture rate for *t_s_*.
- *R_is_*, a random variable representing the average capture efficiency in *i* for all transcripts in *s*.
- *V_is_*, a random variable representing the volume of solution of *s* added to *i*.
- *L_i_*, a random variable representing the cDNA fragment-to-read conversion rate for *i*.
- *l_s_* a constant representing the “sequenceability” of transcripts in *s*.

The product of all of these terms defines the expected number of reads for each *t_s_* in well *i*, and the sum of the products across all *t_s_* is the expected total count of set *s* in *i*. In addition, *ε_is_* represents the effect of sequencing noise on the log-total count, where *E*(*ε_is_*) = 0 and 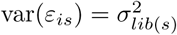.

We assume that *R_is_, V_is_* and *ε_is_* are mutually independent of each other, as they describe separate steps in the protocol. We also assume that *V_is_1__* and *V_is_2__* are independent for sets *s*_1_ and *s*_2_, as each spike-in set is added separately to each well. Similarly, *ε*_*is*_1__ and *ε*_*is*_2__ are assumed to be independent as sequencing noise for each transcript should be unaffected by that of other transcripts. (However, *R*_*is*_1__ and *R*_*is*_2__ are not independent due to well-specific factors affecting capture efficiency for all transcripts). Further details on these variables are provided in Section 1 of the Supplementary Materials.

Let *s* = 1 represent the ERCC spike-in set and *s* = 2 represent the SIRV spike-in set. In the experiment where each spike-in set is added separately to each well, denote the log_2_-ratio of the total counts between the two sets as *θ_i_* = *T*_*i*1_ − *T*_*i*2_ for well *i*. This can also be written as

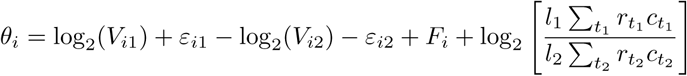
where *F_i_* = log_2_(*R*_*i*1_/*R*_*i*2_) and represents the log-fold change in the average capture efficiency between the two sets (i.e., the difference in behaviour of the transcripts). Computing the variance of *θ_i_* yields

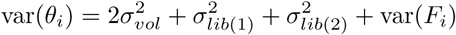
where 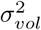 is the variance of both log_2_(*V*_*i*1_) and log_2_(*V_i_*_2_). The volume addition procedure is the same for each spike-in set, so *V_i_*_1_ and *V_i_*_2_ should have the same distribution. We consider the variance of *F_i_* because *R_i_*_1_ and *R_i_*_2_ are not independent (due to well-specific factors, as previously mentioned).

In the experiment where the spike-in sets are premixed before addition, *V_i_*_1_ = *aV_i_*_2_ for some constant *a* representing the proportions in which the two sets are mixed. (This should be close to unity.) If the same premixed solution is added to each well, the relative volume of ERCC spike-ins to SIRV spike-ins must be constant for all wells. This means that the log_2_-ratio for the premixed experiment is

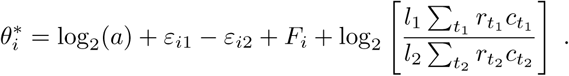

As *a* is constant for all *i*, the variance of 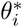 becomes

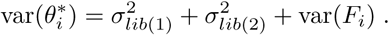

This represents the technical variance attributable to the rest of the scRNA-seq protocol. To obtain an estimate of the variance of the volume addition step, simple arithmetic yields

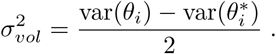

It should be stressed that this variance estimate is relevant to all experiments using the same protocol for spike-in addition, even if the identity or concentration of the spike-in set is different.

With this mathematical framework, we estimated the variance components using the data from our mixture experiments. We observed that the log-ratios *θ_i_* and 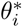 computed from each plate were roughly normally distributed (Supplementary Figure 1). Thus, we fitted a linear model to each set of log-ratios and used the residual variance of the fit as our estimate of var(*θ_i_*) or var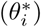. Linear models are particularly useful as they allow blocking on additional structure in the experimental design (Methods). The size of *T_is_* was also similar between wells with premixed or separate addition of spike-ins, which simplifies the calculation of 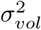 (see Supplementary Figure 2, Section 1 of the Supplementary Materials for details). Finally, the order of spike-in addition did not significantly affect the variance estimates for the separate-addition wells in most plates (Supplementary Figure 3).

Our results indicate that 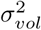 is consistently smaller than the variance in the rest of the protocol (Figure 2a). Indeed, no significant difference was detected between the estimated var(*θ_i_*) and var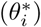 of each plate. This indicates that variability of spike-in volume addition is a minor contributor to the technical variability of the spike-in counts. To put these estimates into context, consider that *T_is_* represents the log_2_-transformed “size factor” for the library generated from well *i*. Spike-in normalization is performed by scaling all counts in this library by the size factor, i.e., 2^−*T_is_*^. This eliminates differences in the coverage of spike-in set *s* between cells and corrects for well/cell-specific technical biases. The variance of the log-size factors is approximately one order of magnitude larger than 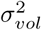 (Figure 2b), which suggests that the latter will not have a major effect on normalization.

**Figure 2.**
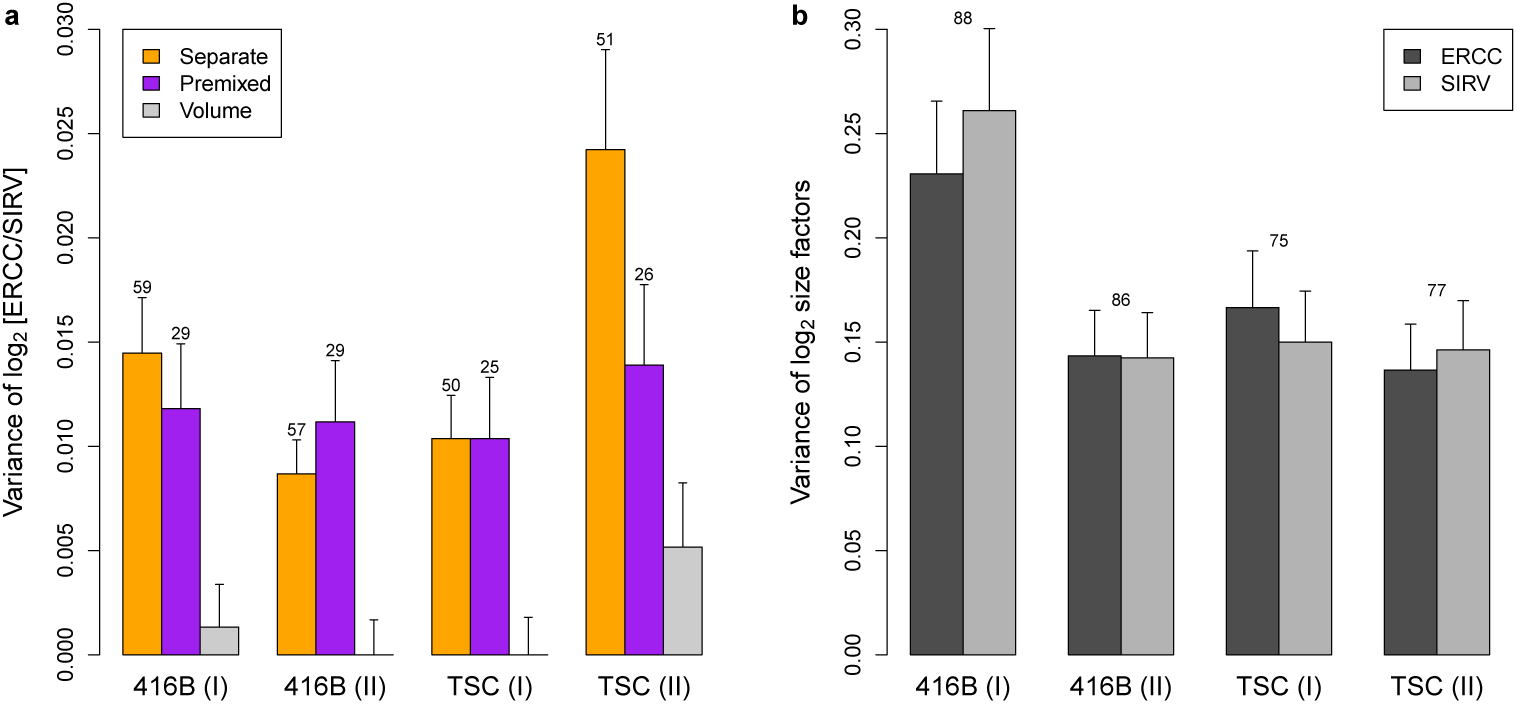
Variance estimates of (a) the log_2_-ratio between the ERCC and SIRV total counts across wells, or (b) the log_2_-size factors computed from those totals. Each estimate is the residual variance of a linear model fitted to the log-ratios or log-size factors across all wells on each plate. Results are shown for experiments with 416B cells or TSCs, with two replicate plates for each cell type. Error bars represent the standard errors of the estimates, assuming log-values are normally distributed. Numbers represent the residual degrees of freedom used for each estimate – for (b), this was the same for each spike-in set. Differences between the separate-addition and premixed estimates for each batch were assessed using a one-sided F-test, yielding *p*-values of 0.28, 1.00, 1.00 and 0.06 from left to right.

### Estimating the variance of differential behaviour

The variance of *F_i_* is also relevant as it determines the effect of differences in behaviour between distinct sets of transcripts. Even when the average capture efficiency differs between sets, spike-in normalization is still appropriate *provided that the fold change in efficiency is the same in all wells*. Consider a situation where there is a consistent increase in efficiency in the spike-in set relative to endogenous transcripts. This scales up the counts for the spike-in transcripts in all libraries by the same amount, which ultimately cancels out between libraries (i.e., the log-fold changes of endogenous or spike-in transcripts between different libraries are unaffected). However, if the fold change in efficiency varies across wells, the accuracy of spike-in normalization is compromised. This is because specific changes in efficiency for the spike-in transcripts are confounded with general changes in efficiency for all transcripts in the well. Differences in the coverage of spike-in transcripts may not represent technical biases affecting other transcripts, precluding their use for normalizing all counts.

In our mathematical framework, the variance of 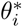 provides an upper bound for the variance of *F_i_*. This quantifies the extent to which normalization is affected by differences in efficiency between two transcript sets. Our estimate of var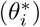 is an order of magnitude lower than the estimated variance of the log-size factors in each plate (Figure 2). This indicates that the potential variance in differential behaviour across wells, while greater than 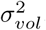, is still relatively small compared to other biases in the system, e.g., differences in cellular RNA content, well-to-well variability in capture efficiency for all transcripts. Here, *F_i_* is computed between two spike-in sets whereas the differences between synthetic spike-in and endogenous transcripts are likely to be greater. Nonetheless, the SIRV and ERCC spike-ins do vary in their biophysical properties (Supplementary Figure 4). For example, the SIRV transcripts have more variable length and lower GC content compared to the ERCC transcripts. This suggests that *F_i_* will include some of the differences in behaviour between synthetic and endogenous RNA, such that var(*F_i_*) can be used as a rough estimate of the magnitude of the associated variability.

We also performed simulations to gauge the relative contribution of var(*F_i_*) and 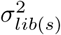 to 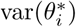 (see Section 2 of the Supplementary Materials). Counts for spike-in transcripts were simulated such that any variability in the log-ratios was only caused by stochastic sampling noise i.e., 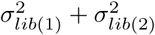. Our results suggest that most of the estimated variance of 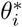 in Figure 2 is driven by sampling noise (Supplementary Figure 5), further reducing the potential impact of differences in behaviour. We also observed that the variance of the log-ratios was robust to decreases in the coverage of the spike-in transcripts in this simulation. In typical experiments, spike-in transcripts take up 5-10% of the library size for each cell (50000-100000 reads in our data). Here, the variance estimates were largely unchanged at 10-fold lower coverage. Thus, spike-in normalization is still reliable when relatively low amounts of spike-in RNA are added or sequenced. This is especially relevant to data sets where the spike-in coverage is lower than recommended, due to difficulties in determining the appropriate concentration of spike-ins to add to each cell when the quantity of endogenous RNA is unknown.

### Assessing the downstream effect of variability with simulations

We assessed whether the results of downstream analyses using spike-in normalization were sensitive to variability in spike-in addition or behaviour. First, we obtained data from plate-based experiments that contained counts for spike-in transcripts. This included public data sets [9, 27] as well as our 416B and TSC data. We then performed analyses such as detection of differentially expressed genes (DEGs) and highly variable genes (HVGs), as well as dimensionality reduction and clustering of cells. This was done without any modification of the data to obtain a set of “original results”.

Next, we designed simulations based on each of the real data sets (see Methods). Briefly, the total spike-in count for each well was rescaled by a randomly sampled factor with variance equal to our experimental estimate of spike-in variance. Counts for the individual spike-in transcipts were rescaled to reflect this new total, thus yielding a simulated data set. Analyses were performed on the simulated data and the new results were compared to the original set of results. Any differences indicate that the analysis is sensitive to spike-in variability in real experiments. The advantage of this simulation design is that only the spike-in counts are modified. Counts for the endogenous transcripts were used directly without any modification, preserving the realistic nature of the data in each simulation.

For DEG detection, we applied edgeR [23] and MAST [5] to the original and simulated data after spike-in normalization. edgeR represents methods designed for DE analyses of bulk RNA-seq data, while MAST represents bespoke single-cell methods. In both cases, we observed only minor (< 5%) changes to the set of significant DEGs upon introducing spike-in variability in each data set (Figure 3a). Similar results were also observed in the top 200 DEGs with the smallest *p*-values, with fewer than 10% of the genes in the set changing across iterations in all scenarios. For HVG detection, we used methods based on the coefficient of variation [3] or the variance of log-expression values [16]. Again, only minor changes were observed in most data sets (Figure 3b), for both the set of significant HVGs and for the top 200 HVGs with the smallest *p*-values. These results suggest that the detection and ranking of DEGs and HVGs are largely robust to variability in spike-in volume or behaviour.

**Figure 3.**
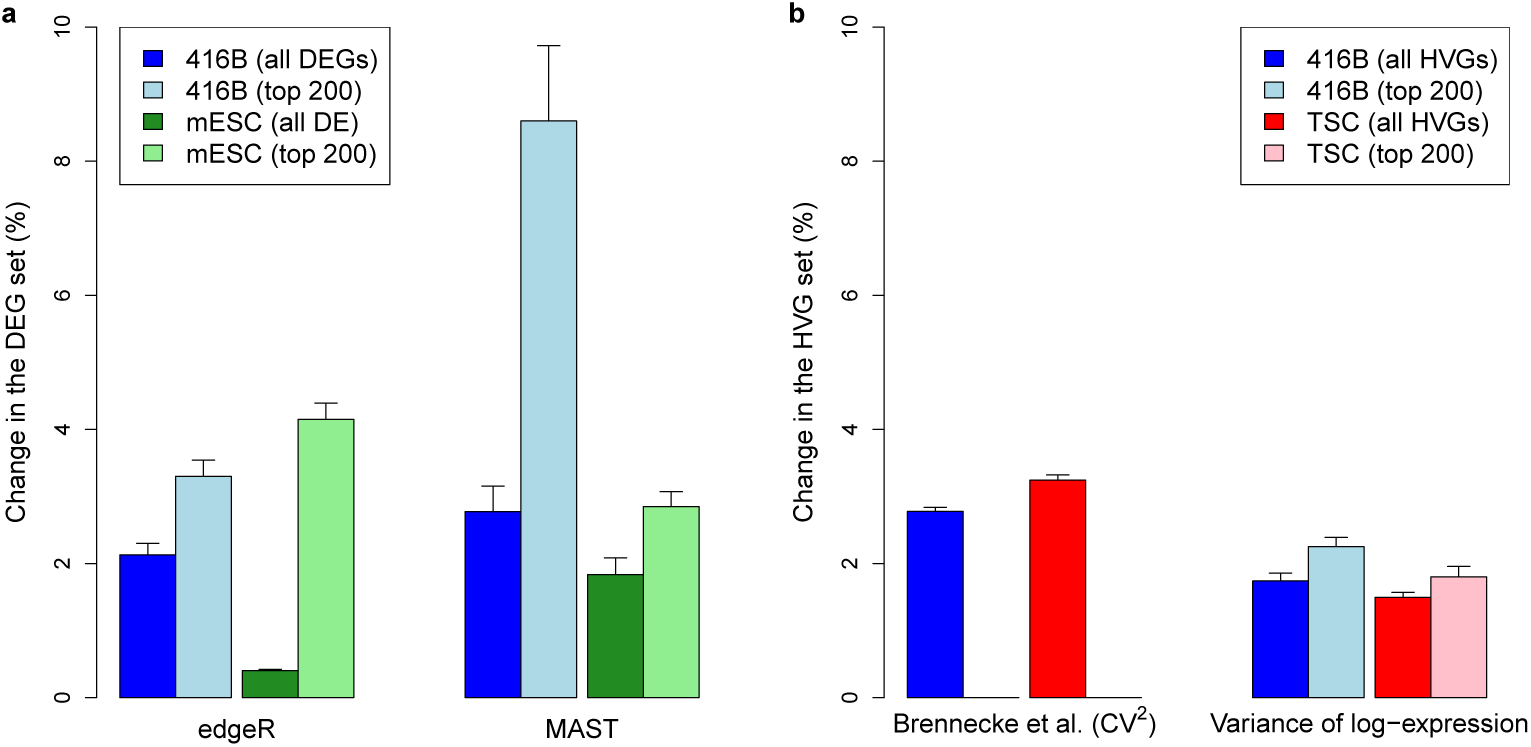
Effect of spike-in variability on DEG or HVG detection in simulated data. (a) The percentage change in the set of DEGs detected in each data set at a FDR of 5% by edgeR or MAST. This was also calculated for the top set of 200 DEGs with the smallest *p*-values. Simulations were based on our 416B data set, to detect DEGs after inducing expression of a *CBFB-MYH11* oncogene compared to a mCherry control (see Methods); or on the data from Islam *et al*. [9], to detect DEGs between mouse embryonic stem cells (mESCs) and fibroblasts. (b) The percentage change in the set of HVGs detected in each data set at a FDR of 5%, using the Brennecke et *al*. method based on the squared coefficient of variation (CV^2^) or with a method based on the variance of log-expression. This was also calculated for the top set of 200 HVGs with the smallest *p*-values. Simulations were based on our 416B and TSC data to detect HVGs across cells. All values represent the mean of 20 simulation iterations, and error bars represent standard errors.

For dimensionality reduction, we restricted ourselves to principal components analysis (PCA) on the normalized expression profiles of all cells. While *t*-distributed stochastic neighbour embedding [31] is commonly used, its robustness is difficult to evaluate due to its randomness. We generated PCA plots of the first three principal components using both the original and simulated data. At each simulation iteration, coordinates of all cells in the simulated plots were mapped onto the corresponding original plots to determine the sensitivity of the original locations to spike-in variability. Figure 4a indicates that changes in the location of each cell across simulation iterations were generally minor. In particular, movement of cells across iterations did not compromise the separation of different cell types. Thus, spike-in variability does not appear to affect the visual interpretation of PCA plots.

**Figure 4.**
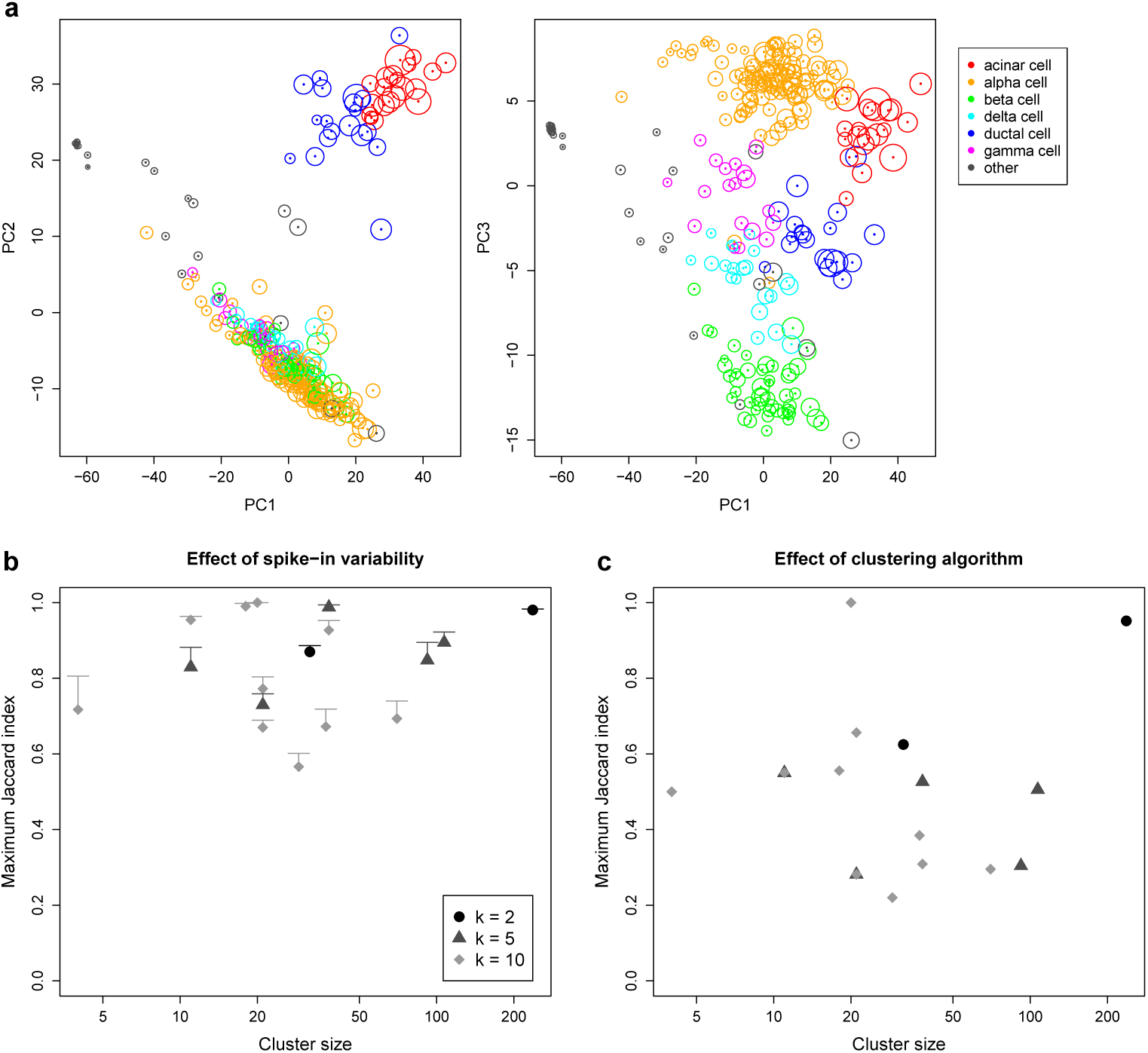
Effect of spike-in variability on dimensionality reduction and clustering in simulated data, based on real scRNA-seq data for cells extracted from a healthy human pancreas [27]. (a) PCA plots of the first three principal components, where each cell is coloured according to its annotated cell type from the original study. The circle around each cell contains 95% of remapped locations across the simulation iterations, and represents the deviation in location due to spike-in variability. (b) Clusters were identified from the original data by hierarchical clustering with Ward’s criterion, followed by a tree cut with *k* of 2, 5 or 10. This was repeated at each simulation iteration, and the maximum Jaccard index between each original cluster and any of the simulated clusters at the same *k* was computed. Each value represents the mean of 20 simulation iterations, and the error bars represent standard errors. (c) The maximum Jaccard index for each original cluster generated with Ward’s criterion compared to the clusters generated from complete-linkage clustering of the original data.

Finally, we performed hierarchical clustering and applied a tree cut to identify clusters of cells in the original data. This was repeated at each simulation iteration to obtain a corresponding set of simulated clusters. For each original cluster, we computed the Jaccard index with respect to each of the simulated clusters and recorded the maximum value across all simulated clusters. A large maximum Jaccard index means that most of the cells in the original cluster are still grouped together in the simulation, i.e., the original cluster is (mostly) successfully recovered in one of the simulated clusters. We observed that the maximum Jaccard indices were moderate to large (Figure 4b), with values above 0.6 for most of the original clusters. To put this into context, we re-clustered the original data using a different algorithm. This yielded smaller Jaccard indices for all clusters (Figure 4c), indicating that spike-in variability has less effect on the results than the choice of clustering method.

## Discussion

In this article, we performed mixture experiments to quantify the variability of spike-in RNA addition across wells in a plate-based scRNA-seq protocol. We also obtained a rough estimate of the well-to-well variability in the differences in behaviour between two different sets of spike-in transcripts. Both values were at least an order of magnitude smaller than the variance of spike-in coverage across cells, suggesting that differences in spike-in volume or behaviour were not major sources of error in the context of spike-in normalization. This was supported by simulations where the introduction of realistic levels of spike-in variance yielded only minor changes in the results of DEG and HVG analyses as well as PCA and clustering. Our results indicate that spike-in normalization is reliable enough for routine use in scRNA-seq data analyses. The common criticisms of using spike-in RNA for normalization are only weakly relevant, if at all, to single-cell transcriptome studies, and can generally be ignored.

Our conclusions differ from those of Risso *et al*. [22], where spike-in normalization is not considered reliable enough for analyses of bulk RNA-seq data. We speculate that this difference may be due to the difficulty of adding an appropriate amount of spike-in RNA at the population level. For example, should spike-in RNA be added at a constant ratio with respect to the concentration of endogenous RNA, or to the number of cells in the sample? If the endogenous RNA concentration or the number of cells determines the amount of spike-in RNA to be added, these will need to be experimentally quantified for each sample. In that case, how accurate is the quantification, and what effect do errors have on the downstream analysis? These questions are not relevant to single-cell experiments where the obvious approach is to add the same amount of spike-in RNA to each individual cell.

We have used the Smart-seq2 protocol in our study to reflect its widespread use in the scRNA-seq literature. However, our estimate of 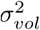 is agnostic to how reverse transcription, amplification and sequencing were performed, as these steps are represented by other mathematical terms. Thus, we expect our conclusions to be broadly applicable to any scRNA-seq protocol where spike-in RNA is added in a similar manner (using repeater pipettes, see Methods). Different results will be obtained using other methods for spike-in addition, e.g., with robotics systems or microfluidics, where volume handling may be even more precise. Our experimental framework may also be useful for evaluating the precision of spike-in addition when developing new scRNA-seq protocols or setting up existing protocols in new laboratories, to ensure that spike-in RNA is added correctly to each cell.

The term var(*F_i_*) represents the variability in the difference in behaviour between the SIRV and ERCC spike-in sets across wells. However, arguably a more relevant quantity is the variability in the difference *P_is_* between synthetic spike-in and endogenous RNA, as this affects the accuracy of normalization. It may be possible to obtain a rough estimate of var(*P_is_*) by using pooled cellular RNA from another organism as one of the spike-in sets [3], so that 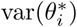 provides an upper bound on the variance in the differences in behaviour between synthetic and endogenous RNA. We chose not to do so because of the difficulty in reproducibly using the same pool of cellular RNA across batches, and in calibrating the concentration of RNA to be added to each well. Use of UMI counts may also provide a tighter bound on var(*F_i_*) or var(*P_is_*) by reducing the contribution of amplification noise to var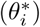.

We stress that our study only examines the reliability of spike-ins for “relative” normalization, i.e., to make counts comparable across cells. We do not consider the reliability of spike-ins for absolute quantification, i.e., to determine the number of molecules of each transcript in each cell. This is more difficult to evaluate as accuracy is affected by the magnitude of the differences in the behaviour of spike-in and endogenous transcripts. In contrast, relative normalization is only affected by variability in the differences in behaviour across wells, as discussed above. Nonetheless, our conclusions are still relevant as absolute quantification depends on the precise addition of spike-in RNA to each cell.

## Methods

### Obtaining and culturing 416B cells and TSCs

The murine multipotent myeloid progenitor cell line 416B [4] was stably transduced with a TetOn construct of the *CBFB-MYH11* (CM) oncogene (type A cDNA), using an in-frame F2A-mCherry protein as a reporter. As a control, cells were alternatively transduced with a version of the construct lacking the CM cDNA. Cells were maintained in RPMI medium, supplemented with 10% fetal calf serum and antibiotics. Expression of the CM oncogene or the mCherry control was induced by treatment with 1 μg/ml of doxycycline, and induction was confirmed after 24 hours by measurement of mCherry levels by fluorescence activated cell sorting (BD Fortessa).

Murine TSCs were kindly provided by Dr. Jennifer Nichols (Wellcome Trust and MRC Cambridge Stem Cell Institute) and cultured by Liliana Antunes (Wellcome Trust Sanger Institute) on mouse embryonic fibroblast (MEF) feeders with TSC culturing medium (a combination of 70% MEF conditioned media (R&D systems) and 30% RPMI 1640, supplemented with 20% FBS, 2 mM L-glutamine, 1 mM sodium pyruvate, 100 μM β-mercaptoethanol, 25 ng/mL human recombinant FGF4 (R&D systems) and 1 μg/mL heparin (Tocris Bioscience)). To prepare for single-cell sorting, cells were harvested with trypsin and MEF feeders were depleted by plating the cells onto a gelatinised plate followed by incubation for 1h at 37°C on TSC culturing medium. The supernatant containing TSCs was used for sorting.

### Spike-in mixture experiments with Smart-seq2

Single-cell RNA sequencing was performed using an adaptation of the previously described Smart-seq2 protocol [20]. Single 416B cells or TSCs were sorted into individual wells of a 96-well microtiter plate. Each well contained 2.3 μl of lysis buffer with RNAse inhibitor (Ambion) in a 0.2% (v/v) Triton X-100 solution. Reverse transcription (RT) was performed in a final volume of 13.2 μl per well, containing 1 μM of oligo-dT (Sigma-Aldrich), 1.04 mM of each dNTP (ThermoFisher), 100 U of SuperScript II retrotranscriptase (Invitrogen/ThermoFisher), 5 U of RNase inhibitor (Ambion), 5 mM of DTT, 1 M of Betaine (Sigma-Alrich), 6 mM of MgCl_2_ (Ambion) and 1 μM of TSO primer (Exiqon). Preamplification was performed in a total volume of 27 μl that contained 13.5 μl of HiFi Hotstart ReadyMix (2×; KAPA Biosystems) and 0.1 μM of IS PCR primer (Sigma-Aldrich). After 23 cycles of amplification, samples were cleaned with 80% (v/v) of Ampure beads (Beckman Coulter). Sequencing libraries were prepared using the Nextera XT DNA sample preparation kit (Illumina). This was repeated to obtain several batches of sequencing data, with each batch consisting of one plate of cells of the same type.

To perform the mixture experiments, spike-in RNA was mixed into the RT reagent solution and added to each well. This was done such that each well contained 0.1 μl of a 1:3,000,000 dilution of the ERCC RNA Spike-In Mix (Invitrogen/ThermoFisher) and 0.12 μl of a 1:3,000,000 dilution of the Spike-in RNA Variant (SIRV) Control Mix E0 (Lexogen). Two separate solutions of RT reagents were prepared for the different spike-in sets. For one third of the wells, addition of the two spike-in sets was performed separately with the RT+ERCC solution first and the RT+SIRV solution second. For another third of the wells, the order was reversed, i.e., with the RT+SIRV solution first and the RT+ERCC solution second. For the remaining wells, the RT+SIRV and RT+ERCC solutions were premixed in a 1:1 ratio and the RT+SIRV+ERCC mixture was added twice to each well. Each addition was performed independently for each well, using a repeater pipette dispensing 2 μl at a time.

Sequencing of the 416B libraries was performed by the Genomics Core facility at the Cancer Research UK Cambridge Institute. The first batch of libraries was sequenced on an Illumina HiSeq 2500 machine generating 125 bp single-end reads, while the second batch was sequenced on an Illumina HiSeq 4000 machine generating 50 bp single-end reads. Sequencing of the TSC libraries was performed at the Wellcome Trust Sanger Institute after library preparation by the Single Cell Genomics Core facility. Both batches were sequenced on an Illumina HiSeq 4000 machine generating 75 bp paired-end reads.

### Data analysis for the mixture experiments

Reads were mapped to the mm10 build of the mouse genome, including sequences of transcripts in the ERCC (https://tools.thermofisher.com/content/sfs/manuals/ERCC92.zip) and SIRV (https://www.lexogen.com/wp-content/uploads/2015/11/SIRV_Sequences_151124.zip) spike-in sets. (The sequence of the *CBFB-MYH11* oncogene was also included in the reference when aligning data from 416B cells.) Mapping was performed using the subread aligner v1.5.1 [13] in RNA-seq mode with unique alignment. The 416B data were aligned in single-end mode while the TSC data were aligned in paired-end mode. Reads with mapping qualities greater than or equal to 10 were assigned to exonic regions of genes using the featureCounts function in the Rsubread package v1.24.1 [14]. Genes were defined using Ensembl v82 annotation for the GRCm38 mouse assembly and annotation for the ERCC and SIRV transcripts. This yielded a count for each endogenous gene and spike-in transcript in each well. Mapping and counting statistics for each batch of libraries are summarized in Supplementary Table 1.

Variance components were estimated from the libraries generated from a single plate. In each well, the sum of counts across all transcripts in each spike-in set was computed, and the log_2_-ratio between the ERCC and SIRV sums was calculated. To estimate var(*θ_i_*), a linear model with a one-way layout was fitted to the log-ratios for all wells where the two spike-in sets were added separately. In each plate of the 416B data set, each combination of treatment (control or oncogene-induced) and spike-in addition order (ERCC or SIRV first) was treated as a group in the one-way layout. In each plate of the TSC data, only the spike-in addition order was used to define the groups. After fitting the model, the mean of the squared residual effects was used as an estimate of var(*θ_i_*). This was repeated for 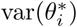 using all wells where premixed spike-ins were added. Here, addition order was irrelevant so the one-way layout contained only the two treatment groups in the 416B data set. Similarly, only a single group was defined for the TSC data. Linear modelling ensures that any changes in the mean log-ratio across groups do not inflate the variance estimate. Note that we fit linear models to each plate separately, to check whether the estimates are consistent across replicate plates.

To detect differences in the variance estimates for premixed and separate addition, an F-test for the equality of variances was applied. Under the null hypothesis of equal variances computed from independent data, the ratio of the variances 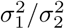 should follow a F-distribution on *n*_1_ and *n*_2_ degrees of freedom, where *n*_1_ and *n*_2_ are the residual degrees of freedom used to estimate 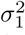 and 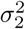, respectively. This can either be one-sided (i.e., 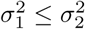 under the null), in which case the lower tail probability at the observed ratio is taken as the *p*-value; or it can be two-sided, in which case the *p*-value is defined as twice the smaller of the two tail probabilities. Significant differences were defined by rejecting the null hypothesis at a type I error rate of 5%. We calculated 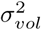 from estimates of var(*θ_i_*) and var 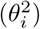 using the expression described above. However, if the difference between var(*θ_i_*) and var 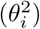 was negative, 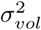 was set to zero instead. To assess the effect of the order of spike-in addition, a linear model was fitted to the subset of relevant wells on each plate to obtain an order-specific variance estimate.

### Simulation design for resampling spike-in variability

For each data set, we compute *T_is_* for each cell *i* and spike-in set *s*. To simplify the design of the simulations, we only consider the ERCC spike-in set here, i.e., *s* = 1. The variance of *T_is_* is

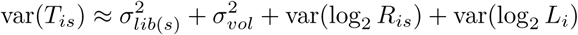

where the approximation assumes that *L_i_* is independent of the other random variables that contribute to *T_is_*. (This is discussed in more detail in Section 1 of the Supplementary Materials.) Let *R_is_* = *R_i_*_0_*P_is_*, where *R_i_*_0_ is the well-specific average capture efficiency of endogenous transcripts and *P_is_* is the fold change in average efficiency of the transcripts in s over their endogenous counterparts. We assume that *R_i_*_0_ and *P_is_* are independent for each well, and that var(log_2_ *P_is_*) can be approximated with var(*F_i_*), i.e., the well-to-well variability in relative capture efficiency between the two spike-in sets is similar to that between spike-ins and endogenous transcripts. This yields

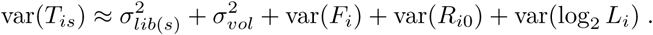

Let us denote 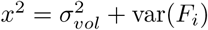, representing the total variance attributable to spike-in addition and capture efficiency. We also denote 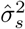 as the estimate of var(*T_is_*) across wells, and 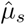 as the estimate of *E*(*T_is_*). We use the estimated 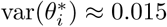 in Figure 2a as our estimate 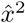 of the upper bound of *x*^2^. This is based on the fact that the estimated 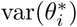 provides an upper bound on var(*F_i_*), while 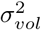 is near-zero in Figure 2a. For each well *i*, we compute a simulated log2-total 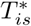 as

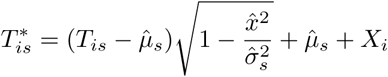
where *X_i_* ∼ Normal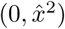 and is independently sampled for each well. This approach ensures that var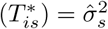. In contrast, if *X_i_* were directly added to *T_is_*, the variance of 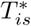 would be inflated as *x^2^* is already present in var(*T_is_*), i.e., the contribution of spike-in variance would be doubled.

Counts for the library generated from each well were rescaled to reflect the new, simulated log-total. A quantile adjustment approach was used to preserve the empirical mean-variance relationship. Briefly, a negative binomial generalized linear model (NB GLM) was fitted to the counts across all wells for each spike-in transcript, using the mglmOneGroup function in edgeR [18, 23] with an all-intercept design matrix and *T_is_* (converted to base *e*) as the offset for well *i*. An abundance-dependent trend was also fitted to the NB dispersions across all spike-in transcripts using the estimateDisp function. For each transcript *t*, we assumed that the count *y_ti_* for well *i* was sampled from a NB distribution with mean equal to the corresponding fitted value of the GLM and dispersion equal to the fitted value of the mean-dispersion trend. We scaled the NB mean by 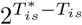 to obtain a modified NB distribution. Using the q2qnbinom function [25], we calculated the lower tail probability of *y_ti_* in the original distribution and identified the corresponding quantile with the same tail probability in the modified distribution. This new quantile was used as the simulated count for transcript *t* in *i*.

### Evaluating the robustness of DEG detection

Two data sets were used to test the effect of spike-in variability on DEG detection. The first was the 416B data generated previously, where DEGs were detected between control and oncogene-induced cells in both plates. Here, we used an additive model with a treatment term and a blocking factor for the plate. The second data set was obtained from the NCBI Gene Expression Omnibus (GEO) with the accession number GSE29087, and compared mouse embryonic stem cells and fibroblasts [9].

In both studies, DEGs were detected between conditions using edgeR and MAST. Implementation details of each method are provided in Section 3 in the Supplementary Materials. Briefly, normalization was performed by scaling the counts (explicitly or via offsets) such that the spike-in totals were the same between cells. The set of DEGs in the original data was then identified at a FDR of 5%. This procedure was repeated for the simulated data, and the number of genes that were detected in the original results and not in the simulated results (or vice versa) was recorded as a proportion of the total number of original DEGs. The proportion of the top 200 genes with the smallest *p*-values that were shared between the original and simulated results was also computed. This was repeated for 20 simulation iterations and the average proportion across iterations was reported for each method.

### Evaluating the robustness of HVG detection

Our 416B and TSC data sets were used to assess the effect of spike-in variability on detection of HVGs. In the former, blocking was performed to remove plate- and treatment-specific effects on mean expression, i.e., HVGs were detected within treatment conditions on each plate. Similarly, blocking was performed on the plate of origin for each cell in the TSC data set to remove plate effects.

In each data set, spike-in normalization was performed and HVGs were detected using two approaches based on spike-in counts (See Section 3 in the Supplementary Materials for implementation details of each method.) The first approach is based on the method of Brennecke *et al*. [3] where the squared coefficient of variation for each gene is tested for a significant increase above technical noise. The second approach is based on the variance of the log-normalized expression values [16], which provides some more robustness against outlier expression patterns. Each method was applied on the original and simulated data, and a set of significant HVGs was detected at a FDR of 5%. The proportion of HVGs common to both the original and simulated sets was computed, along with the common proportion among the top 200 genes with the lowest *p*-values. This was repeated for 20 simulation iterations and the average proportion across iterations was reported for each method.

### Evaluating dimensionality reduction and clustering

Count data from a study of pancreatic islet cells [27] were obtained from ArrayExpress with the acession E-MTAB-5061. Spike-in normalization was performed and a set of HVGs was defined using the variance-of-log-expression method. PCA plots of the first three components were constructed from the matrix of log-expression values for the HVGs. This process – including HVG detection – was repeated with the simulated data after introducing spike-in variability. To compare each simulated PCA plot to the original plot, the coordinates of each cell in the former were mapped onto the latter by rescaling and rotation. Robustness was assessed based on the spread of remapped coordinates across all simulation iterations for each cell. See Section 3 in the Supplementary Materials for details.

To test the robustness of clustering, the matrix of Euclidean distances between cells was computed from the HVG log-expression values. Hierarchical clustering was performed using the Ward criterion and the resulting dendrogram was cut into 2, 5 or 10 clusters. (This was done using the hclust and cutree commands, respectively, from the stats package.) This process was repeated with the simulated data, and the Jaccard index between every pair of simulated and original clusters was computed. For each original cluster, the maximum Jaccard index across all simulated clusters was recorded at each simulation iteration. This value represents the extent to which the membership of the original cluster was preserved in the most similar simulated cluster. We also compared the original clusters to those generated from complete-linkage clustering of the original HVG log-expression values.

## Data access

Data are available on ArrayExpress using the accession E-MAT-5522. Code used for the statistical analysis and simulations are available at https://github.com/MarioniLab/SpikeIns2016.

## Acknowledgements

We thank Jennifer Nichols and Liliana Antunes for supplying the TSCs. We also thank Victoria Moignard and Wajid Jawaid for helpful discussions about the experimental design.

## Author contributions

ATLL proposed the mixture experiments and performed the statistical analysis and simulations. FJCN adapted the Smart-seq2 protocol for the spike-in mixtures and generated the 416B data. LHV generated the TSC data. BG and JCM provided direction and guidance on the project. All authors wrote and approved the manuscript.

## Supplementary materials

The Supplementary Materials is a single PDF file that consists of Sections 1-3 and contains Supplementary Figures 1-5 and Supplementary Table 1.

